# Restriction Point timing and cell cycle variability – a re-evaluation

**DOI:** 10.1101/2020.07.09.186700

**Authors:** Robert F. Brooks

## Abstract

The Restriction Point (R) in the mammalian cell cycle is regarded as a critical transition in G1 when cells become committed to enter S phase even in the absence of further growth factor stimulation. Classic time-lapse studies by Zetterberg and Larsson suggested that the acquisition of growth factor independence (i.e. passage of R) occurred very abruptly 3-4 hours after mitosis, with most cell cycle variability arising between R and entry into S phase. However, the cycle times of the post-R cells that continued on to mitosis after serum step-down without perturbation were far less variable than the control cells with which they were compared. A re-analysis of the data, presented here, shows that when the timing of R and entry in mitosis are compared for the same experiments, the curves are superimposable and statistically indistinguishable. This indicates that the data are compatible with the timing of R contributing to much of the overall variability in the cell cycle, contrary to the conclusions of Zetterberg and colleagues.

## Introduction

The recent developments in live-cell imaging of cell cycle markers have brought renewed focus on cell cycle variability, with extensive data showing that ostensibly identical cells undergo critical transitions at widely differing cell ages [1–12]. Although such variability has long been recognised [13;14], the molecular basis for it remains poorly understood. Inheritance of DNA damage from the preceding cycle may contribute to some of the variability [1;3;11]. However, with proliferating Hs68-TERT human fibroblasts, for example, only 10-15% of cells display DNA-damage foci, indicating that for the majority of cells, the variability has some other cause [15].

For mammalian cells, the Restriction Point (R) in G1 is widely regarded as a critical step in the cell cycle when cells become committed to enter S phase and subsequently divide, independent of further growth factor stimulation [16–20]. Prior to R, removal of growth factors leads to exit from the cell cycle into the G0 quiescent state; after R, cells continue into S phase and mitosis even after growth factor withdrawal. For normal, growth factor-dependent cells, R is widely regarded as an important decision point that must be passed in each and every cell cycle [16–20].

In any attempt to understand the origin of cell cycle variability, establishing the timing of R in relation to the onset of S phase is clearly important. By following exponentially-growing 3T3 cells using time-lapse microscopy, Zetterberg and Larsson concluded that growth factor independence was acquired very abruptly, 3-4 hours after mitosis and some time before entry into S phase [21]. Thus, in response to serum-removal, almost all cells younger than 3-4 hours underwent growth arrest whereas all cells older than this continued on to mitosis with no delay or set-back. Similar results were reported for proliferating normal human fibroblasts and mammary epithelial cells [22;23]. The acquisition of growth factor independence (passage of R) appeared to be much more abrupt than the subsequent entry into S phase or mitosis, leading to the suggestion that practically all of the variability in the cell cycle arises *after* R, in the interval prior to the onset of S phase, which they called G1ps [21;22;24]. However, these observations for exponentially growing cells are in contrast to results for quiescent 3T3 cells responding to serum stimulation where growth factor independence was acquired at variable times relative to step-up, but at a roughly fixed interval (of around 5 hours) before entry into S [25], consistent with the variability arising *before* R. Similarly, recent studies of non-established human fibroblasts also concluded that the timing of R was variable rather than fixed [10]. Indeed, for some cell types, many cells fail to undergo immediate cell cycle arrest after growth factor withdrawal in early G1, suggesting that in these cases, passage of R had occurred before mitosis, in the mother cell [9;10;12].

In view of the uncertainty as to the timing of R relative to entry into S phase, the 3T3 data from Zetterberg and Larsson [21] have been re-examined here. This reanalysis does not support the idea that most cell cycle variability arises after R. Rather, the data are compatible with the timing of growth factor independence (passage of R) contributing to much of the variability in cycle times.

## Results and Discussion

The time-lapse data from Zetterberg and Larsson [21] showing the consequences of transient serum-removal as a function of cell age, are reproduced in Fig. 1. All cells older than 4 hours showed no perturbation and divided on schedule. For cells younger than 4 hours, even a 30 minute serum withdrawal (Fig. 1A) was sufficient to delay the cell cycle of many cells. When serum was removed for 1 hour or more (Fig. 1B, C), all cells younger than 3 hours in these experiments experienced an excess delay of 8 hours [21] over and above the interval of serum deprivation, consistent with a set-back into G0. Between the ages of 3 to 4 hours, the fraction of cells that had passed R (i.e. became independent of serum and continued on to mitosis on schedule) increased very steeply (curve a, Fig. 2), much more steeply than for entry into S phase (curve b, Fig. 2) or mitosis (curve c, Fig. 2). Such observations led Zetterberg and colleagues to conclude that practically all the variability in the cell cycle arises in G1 after passage of R (acquisition of growth factor independence) [24]. However, inspection of Fig. 1 shows that among the cohort of cells unperturbed by serum step-down (i.e. those older than 4 h) there are no cycle times less than 13.5 h or greater than 20 h. This suggests a much narrower distribution of cycle times than in the control cells (Fig. 2, curve c), which ranges from around 10 h to more than 24 h. To investigate this further, the cycle times of all cells older than 4 h were extracted from Fig. 1 (A-C) and the cumulative distribution for entry into mitosis determined. This is shown superimposed on the Zetterberg and Larsson data as curve d in Fig. 2. Curve d is clearly much steeper than curve c and the two distributions are significantly different (χ^2^ = 87.7, 8 df, p<0.001, using an estimated sample size of 1000 for the control population based on the stated analysis of 100-300 cells in each of 5 experiments [21]; for a sample of 1500, the value of χ^2^ is even larger). Evidently, the unperturbed cell cycles of post-R cells in Fig. 1 are much less variable than the control cells in Fig. 2, curve c, suggesting either that they are drawn from entirely different populations or that the control data, pooled from 5 separate experiments [21], includes significant inter-experimental variability not seen in Fig. 1. Either way, it is inappropriate for the data in Fig. 1 to be compared directly with the “control” cells in Fig. 2 (curve c).

**Fig. 1.**
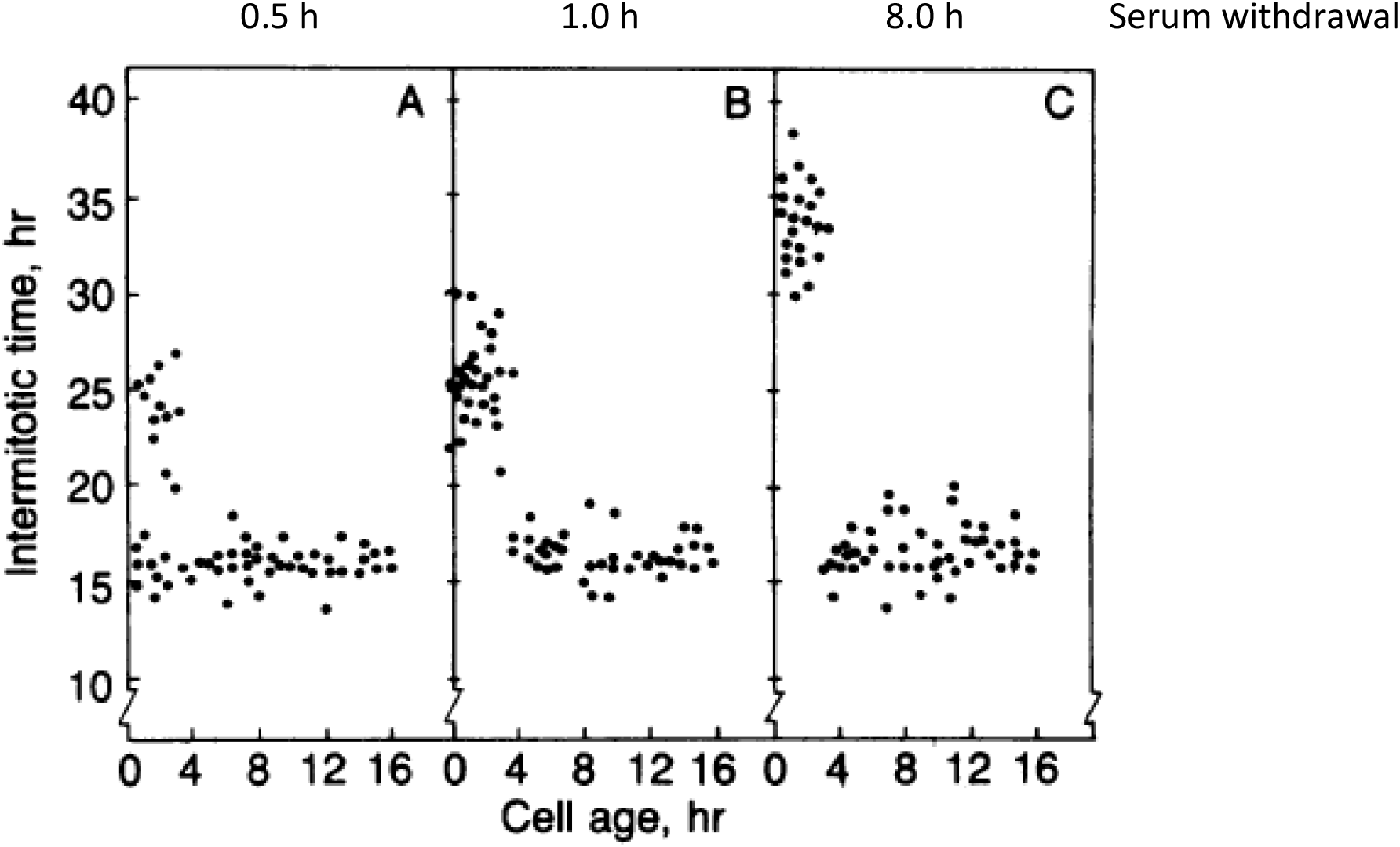
The relationship between cell age at the time of serum deprivation and subsequent intermitotic time, from time-lapse microscopy (reproduced from Fig. 1 of Zetterberg and Larsson[21]). Cells were transferred transiently to serum free medium for 0.5 h (A), 1.0 h (B) or 8 h (C).

**Fig. 2.**
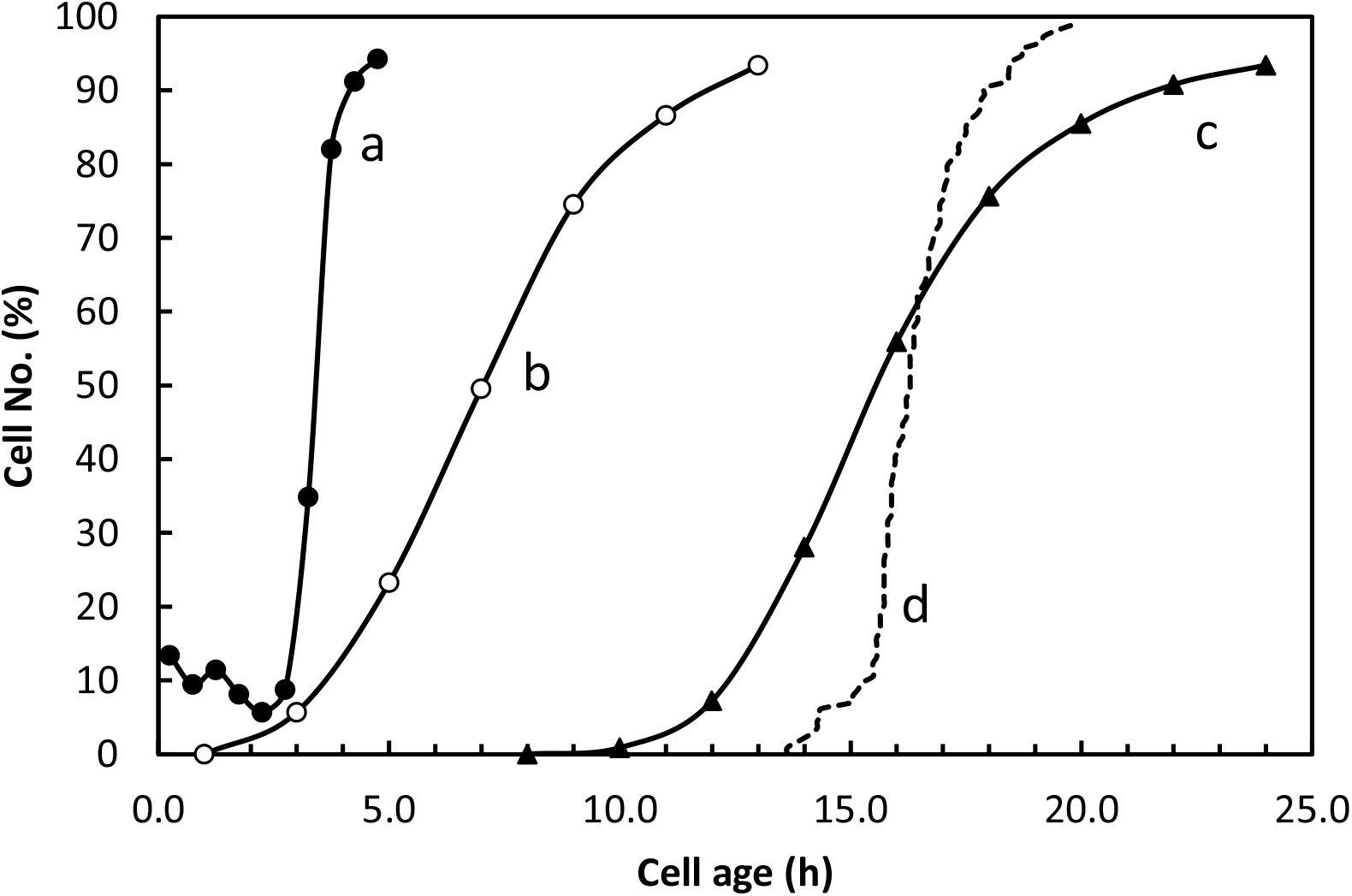
Cell age distribution for different cycle phases. Curves a, b and c are redrawn from Fig. 3 of Zetterberg & Larsson [21]. **Curve a** (closed circles): passage of R (acquisition of serum independence) – the “G1 pm” curve of Zetterberg & Larsson. **Curve b** (open circles): entry into S phase in proliferating control cells (proportion of cells labelled with [^3^H]-thymidine as a function of cell age). **Curve c** (closed triangles): entry into mitosis for proliferating control cells. **Curve d** (dashed line): entry into mitosis for the unperturbed cohort of cells (i.e. > 4 h old) in Fig. 1, A-C (N=114 cells).

It is noteworthy that the rate of entry into mitosis for the unperturbed cells of Fig. 1 (curve d in Fig. 2) is very similar to the timing of growth factor independence (curve a in Fig. 2). To compare these more carefully, the difference in the median values (12.85 h) for the two curves was subtracted from the times of mitosis, so as to superimpose the distributions (Fig. 3A). In addition, an estimate of the 95% confidence intervals for the time points for growth factor independence was included, assuming a sample size of 40 cells in each of the 10 class intervals (of 30 mins) spanning the cells aged 0-5 h. This is an overestimate based on a maximum of 1500 cells analysed (100-300 cells in each of 5 experiments [21]) of which roughly 25% would have been in the 0-5 h age range (^~^375 cells), giving ^~^37 in each of the ten 30 minute class intervals. In practice, the number of cells in each 30 minute class interval is likely to have been even smaller, meaning that the actual confidence intervals are even broader than shown. As may be seen in Fig. 3A, the curves for growth factor independence (passage of R) and entry into mitosis are remarkably similar. After the 50% values it is possible that the rate of entry into mitosis may slow slightly compared to passage of R. However, the data points for entry into mitosis lie within (or are very close) to the 95% confidence intervals for the timing of growth factor independence. There is therefore little evidence for any additional variability arising between R and entry into mitosis, contrary to the conclusions of Zetterberg and Larsson [21].

**Fig. 3.**
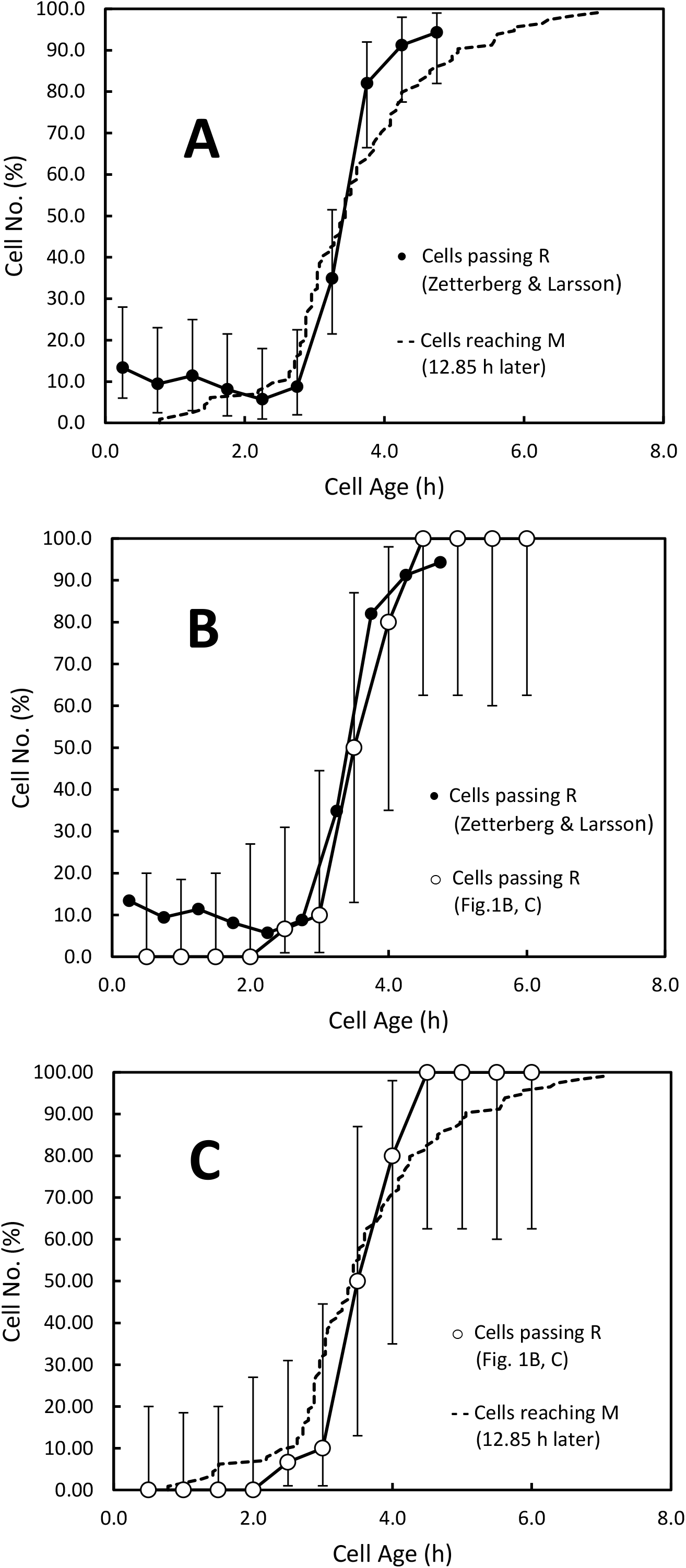
Timing of serum-independence (passage of R) and entry in mitosis. **(A)** A comparison of the original data of Zetterberg & Larsson [21] for the timing of serum independence with entry into mitosis for the unperturbed (post-R) cells of Fig. 1 B, C. **(B)** A comparison of the original data of Zetterberg & Larsson [21] for the timing of serum independence with a similar analysis for the cells in Fig. 1 (B, C). **(C)** A comparison of entry into mitosis with the timing of serum independence for cells in the same experiment (Fig. 1) **(A, C)** *<u>Dashed line</u>*: Cumulative distribution for entry into mitosis for the unperturbed cells (>4 h old) in Fig. 1A-C, after subtracting 12.85 h from the intermitotic times (the difference in median values for curves a and d in Fig. 2). **(A, B)** *<u>Closed circles</u>*: passage of R (acquisition of serum independence) – same as curve a in Fig. 2 – data from Fig. 3 of Zetterberg & Larsson [21]. Also included in (**A**) are the 95% confidence intervals for proportions (from Table 41 of the Biometrika Tables for Statisticians Vol. 1 [32]) for an estimated sample size of 40 cells in each 30-minute age class (see text). **(B, C)** *<u>Open circles</u>*: passage of R (acquisition of serum independence) for the cells in Fig. 1 (B, C) together with the 95% confidence intervals for proportions (from Table 41 of the Biometrika Tables for Statisticians Vol. 1 [32]). For sample sizes, see Table 1. The points are the % of cells that had become serum-independent (post-R), for successive hourly cohorts, staggered by 30 minutes (i.e. cells aged 0-1, 0.5-1.5, 1-2, 1.5-2.5, etc.), plotted at the midpoint of the class, as in Brooks et al [27].

In Fig. 3A, the times of mitosis are drawn from a restricted set of experiments (specifically, those in Fig. 1), whereas the timing of R is based on data pooled from a larger number of experiments (5, see [21]). It could be argued therefore that the two distributions (Fig. 3A) should not be compared directly as they are not drawn from precisely the same population. Accordingly, the timing of R (growth factor independence) was determined for the experiments in Figs. 1B, C that contributed to the estimate of intermitotic times. (Data from Fig. 1A were not included in the timing of R as the period of serum withdrawal – just 30 mins – was too short to delay the cell cycle of many cells younger than 4 h.) A limitation of this analysis, inherent to the experimental approach, is the very small sample size. The number of cells in each age cohort is very small indeed even when grouped into class intervals of 1 h (Table 1) instead of 30 minutes as in [21]. As a result, the 95% confidence intervals are very broad. Despite this, it is clear that the timing of R for the cells of Figs. 1B, C is indistinguishable from that for the pooled data (Fig. 3B). In addition, the 95% confidence intervals for the timing of R overlap completely with the timing of mitosis (minus 12.85 h) within the same set of data, eliminating any concerns over inter-experimental variation (Fig. 3C).

**Table 1.** Numerical data for the acquisition of serum independence (passage of R) for the cells of Fig. 1 (B, C), corresponding to the open circles in Fig. 3, to indicate sample sizes. Post-R cells are those that pass on to mitosis without perturbation after serum withdrawal.

| **Cell age (h)** | **Total No. of cells** | **No. post-R** | **% post-R** |
| --- | --- | --- | --- |
| 0 – 1.0 | 17 | 0 | 0.0 |
| 0.5 – 1.5 | 18 | 0 | 0.0 |
| 1.0 – 2.00 | 17 | 0 | 0.0 |
| 1.5 – 2.5 | 12 | 0 | 0.0 |
| 2.0 – 3.0 | 15 | 1 | 6.7 |
| 2.5 – 3.5 | 10 | 1 | 10.0 |
| 3.0- 4.0 | 4 | 2 | 50.0 |
| 3.5 – 4.5 | 5 | 4 | 80.0 |
| 4.0- 5.0 | 8 | 8 | 100.0 |
| 4.5 – 5.5 | 8 | 8 | 100.0 |
| 5.0 – 6.0 | 7 | 7 | 100.0 |
| 5.5 – 6.5 | 8 | 8 | 100.0 |

The studies from Zetterberg and colleagues [21–24] are an important contribution towards understanding the origin and location of variability in the cell cycle, and the single-cell, time-lapse analysis used by them is very powerful. Nevertheless, the approach suffers from certain limitations. Firstly, in any time-lapse study, some cells are lost to view owing to migration from the field of view or are lost to further analysis by the end of the film. Because the probability of being lost increases with the length of time a cell has been followed, lost cells must be included in any analysis up to the point they are lost, to avoid bias [26]. In the studies of Zetterberg and colleagues [21–23], this does not seem to have been done. As a result, there is likely to be a deficiency of cells with long cycle times in the data collected, which will affect the shape of the distributions presented, possibly accounting for the differences between the control cells and those subjected to serum withdrawal. Secondly, there is the issue of small sample sizes. The number of cells in each narrow age-range, at the time of serum step-down, is tiny (Table 1; see also [27] for a similar analysis). Estimates of the proportion of cells in each age cohort that have or have not become independent of serum growth factors are therefore subject to considerable statistical uncertainty, even after pooling data from multiple experiments to increase sample size. Such pooling carries the risk of introducing additional variability due to the almost inevitable differences between experiments. This could be a reason for the much greater cell cycle variability in the (pooled) control population than in the unperturbed cohort (cells >4 h old, Fig. 1), c.f. curves c and d Fig. 2. When instead the timing of growth factor independence is compared directly with the intermitotic times of the unperturbed cohort for the same experiments, the distributions are superimposable and statistically indistinguishable (Fig. 3c). The re-analysis presented here is therefore compatible with most cell cycle variability originating at or before the point of growth factor independence and not after it as previously suggested [24].

Lastly, it is worth noting that conventional understanding of the Restriction point may require revision [20]. It is widely accepted that R involves hyperphosphorylation of the retinoblastoma protein (RB) by CDKs to release E2F family transcription factors that drive gene expression needed for cell cycle entry [17–20;28]. Growth factor stimulation leads to the expression of cyclin D and the activation of CDK4,6 which in turn initiates phosphorylation of RB. The E2F released leads to expression of cyclin E, and later cyclin A, activating CDK2, further driving phosphorylation of RB in a positive feedback loop [17;18;28]. Passage of R is generally equated with a point when RB hyperphosphorylation passes from being cyclin D (and growth factor) dependent to cyclin E/A dependent (and insensitive to growth factor removal) [17;18;28]. However, recent data from Chung et al [6] show that RB hyperphosphorylation remains dependent on Cyclin D-CDK4,6 (as determined by sensitivity to the CDK4,6 inhibitor palbociclib) right up to the very end of G1, some time after the point of growth factor independence. Only near the end of G1 does RB hyperphosphorylation become driven solely by cyclin E/A-CDK2, without the need for Cyclin D-CDK4,6 [6]. This coincides with the point in the cell cycle when the ubiquitin ligase APC-CDH1 is inactivated and proteolysis shifts from being APC-CDH1 mediated to SCF dependent, only after which cyclin A and other regulators of S phase are able to accumulate [6]. This, it has been suggested, may be the true point of no return in the cell cycle [4;5]. That it occurs several hours after the acquisition of growth factor independence is because cells retain a physiological memory of growth factor stimulation for some time after the external growth factors are removed [6;11]. In particular, although the level of cyclin D falls after growth factor withdrawal, in many cells it remains sufficiently high for long enough, for the cell to pass on to the point when CDK2-cyclin E/A is able to maintain RB hyperphosphorylation without further need of cyclin D-CDK4,6 [6]. The Restriction Point is therefore not an obligatory decision point that must be passed in each and every cell cycle, but a physiological state characterised by a threshold level of Cyclin D-CDK4,6 activity (or CDK2-cyclin E in turn), above which cells are able to continue on to the cell cycle commitment point (probably APC-CDH-1 inactivation) in the absence of further growth factor stimulation. On that basis, asynchrony in the timing of R is merely a reflection of the heterogeneity in Cyclin D-CDK4,6 levels (or Cyclin E-CDK2) at the time of growth factor step-down [6;11]. Thus, cells born with a level of Cyclin D-CDK4 activity above the threshold (leading in turn to a rising level of Cyclin E-CDK2 activity) would pass straight on through the cell cycle in the absence of growth factor stimulation as though they had already passed R before birth [7;9–12]. Such cells would correspond to those where cyclin E/A-CDK2 activity rises immediately after birth, even after mitogen withdrawal [7;9;11;12]. Heterogeneity in levels of Cyclin D-CDK4,6 activity could also account for the differences between cells in growth factor sensitivity, even within a clone, such that some cells continue cycling through multiple cycles in limiting growth factor concentrations while others exit to quiescence [9;14;29–31]. If so, then the reasons for the heterogeneity in Cyclin D-CDK4,6 levels in cells of the same age are of particular importance.

## Methods

### Data extraction and analysis

Figs 1 and 3 of Zetterberg & Larsson (1985) [21] were enlarged and printed at maximum size on A3 paper. The values of the data points were then extracted by direct measurement, using a ruler.

In Fig. 1 of Zetterberg and Larsson (reproduced here as Fig. 1), no cells older than 4 hours at the time of serum withdrawal experienced a cell cycle delay and so were deemed to have passed R (the point of mitogen independence). The cumulative distribution of times to mitosis for these “unperturbed” cells, from all three panels in Fig. 1, was determined using Excel, the sample size being 114. In order to compare this distribution with that of the proliferating control cells of Zetterberg and Larsson (Fig. 2, curve c) using the χ^2^ test, it was necessary to assume a value for the sample size of the latter since this was not stated explicitly. Instead, the data were said to be derived from the analysis of 100-300 cells in each of 5 experiments, indicating a sample size of between 500 and 1500. Accordingly, a value of 1000 was used here, though taking other values at the extremes of the range made little difference to the estimate of χ^2^. The class intervals used in the calculation of χ^2^ were governed by the data points in Fig. 2, curve c, of Zetterberg and Larsson, namely 8-10, 10-12, 12-14, …….., 22-24, >24 h.

To determine the time-course for passage of R, cells in Fig. 1B, C were ranked according to age and grouped into rolling 1 h cohorts staggered by 30 minutes, i.e. 0-1, 0.5-1.5, 1-2, 1.5-2.5, etc.. (Panel A of Fig. 1 was not used here as the period of serum withdrawal – 30 minutes – was too short to delay the cell cycles of many cells in the 0-4 h range.) The number of cells in each 1 h cohort is given in Table 1. The cells in each cohort were then categorised as either pre- or post-R. A cell was considered to be pre-R if its cell cycle was delayed by serum withdrawal compared to the “unperturbed” cells (those older than 4 h at the time of serum withdrawal). For Fig. 1C (serum withdrawal for 8 h), the cells experiencing a delay are clearly separated, as a group, from the unperturbed fraction, and there is no possible ambiguity. For Fig. 1B (serum withdrawal for 1 h only), the delayed group are closer to the unperturbed group. In this case, a cell was considered to have been delayed if its cell cycle was at least 3 h longer than the longest cycle of the unperturbed group (those older than 4 h at serum withdrawal). The fraction of post-R cells in each of the hourly cohorts was then determined (Table 1) and the 95% confidence intervals for proportions estimated by interpolation from Table 41 of the Biometrika Tables for Statisticians Vol. 1 [32].

## Acknowledgements

The author thanks A Zetterberg and O. Larsson for permission to reproduce Fig. 1 and redraw data in Fig. 2.

